# PRMT5 links lipid metabolism to contractile function of skeletal muscles

**DOI:** 10.1101/2022.11.04.515165

**Authors:** Kun Ho Kim, Zhihao Jia, Madigan M. Snyder, Jingjuan Chen, Jiamin Qiu, Stephanie N. Oprescu, Xiyue Chen, Sabriya A Syed, Feng Yue, Bruno T. Roseguini, Anthony N. Imbalzano, Changdeng Hu, Shihuan Kuang

## Abstract

The skeletal muscle plays a key role in systemic energy homeostasis besides its canonical contractile function, but what couples these functions is poorly defined. Protein Arginine MethylTransferase 5 (PRMT5) is a well-known oncoprotein but also expressed in healthy tissues with unclear physiological functions. As adult muscles express high levels of *Prmt5*, we generated myocyte-specific *Prmt5* knockout (*Prmt5^MKO^*) mice. We observed reduced muscle mass, oxidative capacity, force production and exercise performance in *Prmt5^MKO^* mice. The motor deficiency is associated with scarce lipid droplets in myofibers due to defects in lipid biosynthesis and degradation. First, *Prmt5^MKO^* reduced demethylation and stability of Sterol Regulatory Element-Binding Transcription Factor 1a (SREBP1a), a master regulator of *de novo* lipogenesis. Second, *Prmt5^MKO^* impaired the repressive H4R3Me2s (histone H4 arginine-3 symmetric demethylation) at the *Pnpla2* gene, elevating the level of its encoded protein ATGL, the rate-limiting enzyme catalyzing lipolysis. Accordingly, myocyte-specific double knockout of *Pnpla2* and *Prmt5* normalized muscle mass and function. Together, our findings delineate a physiological function of PRMT5 in linking lipid metabolism to contractile function of myofibers.

## Introduction

Skeletal muscles cells (also called myofibers or myocytes) are multinucleated contractile units that empower body movements and mobility. The skeletal muscle also plays a key role in systemic energy homeostasis through glucose disposal and fatty acid oxidation (FAO) ^1,2^. A progressive decline in skeletal muscle mass and function not only impairs exercise capacity but also increases the risks of metabolic disorders associated with cellular oxidative stress, mitochondrial dysfunction, and insulin resistance^3^. Muscle atrophy, one of the most destructive features of muscular dysfunction, results from protein degradation that reduces muscle mass. Additionally, an oxidative-to-glycolytic conversion of myofibers renders them more prone to protein degradation, leading to nutrient-related atrophy^4–6^’.

Intramyocellular lipid (IMCL) is a crucial energy source for FAO, serves as an important building block for cellular and organelle membranes, and generates bioactive metabolites that mediate various signaling pathways^7,8^. IMCL is typically stored in lipid droplets (LDs), cytoplasmic organelles that store and release lipids such as triglycerides (TAG). Dynamics of LDs are linked to a myriad of physiological and pathological processes^9–11^. LD biogenesis is mediated by TAG-synthesizing enzymes under the control of transcriptional factors such as SREBP1^12^. LD turnover is mediated by lipolysis that liberates fatty acids (FAs) from TAG for oxidation as a source of energy. The first and rate-limiting step in lipolysis is catalyzed by adipose triglyceride lipase (ATGL, encoded by *Pnpla2* gene) to hydrolyze TAG to generate a free FAs and DAG (diacylglycerols)^13,14^. Dysfunctional ATGL-mediated lipolysis in myocytes reduces the availability of FAs for energy conversion and impairs muscle function^13,15–17^. Conversely, increased ATGL activity is associated with muscle wasting in cancer cachexia^18^. Despite the known function of ATGL in skeletal muscle and muscle satellite cells^19–22^, what regulates lipid metabolism in the skeletal muscle is poorly understood.

Protein methylation, a common post-translational modification (PTM) that results in the addition of methyl groups to lysine or arginine residues, has been involved in protein functional modulation, subsequent alteration of signaling transduction and gene transcription^23–26^. Protein arginine methyltransferases (PRMTs), composed of nine paralogs, can catalyze methylation of a broad range of substrates, including histone proteins^23,27^. Histone arginine methylation is associated with dynamic gene regulation, as chromatin structure remodeling either leads to activation or repression of targeted genes ^28,29^. For example, symmetric dimethylation of histone 4 arginine 3 (H4R3Me2s) associated with several genes has been reported to underly pathogenesis of cancer ^30–32^. Recent studies have reported that PRMT1, PRMT4, PRMT7 regulates a variety of biological processes in the skeletal muscle^33–36^. Previous studies also reported that PRMT5 is indispensable for the proliferation and differentiation of myogenic progenitors and skeletal muscle regeneration^37,38^.

In this study, we characterized the function of PRMT5 in myocytes through *Myl1^cre^*-driven knockout of *Prmt5* (*Prmt5^MKO^*) in mice. The *Prmt5^MKO^* mice exhibited reduced muscle mass and body weight along with impaired contractile function and motor performance. The *Prmt5^MKO^* mice also contained an increased proportion of glycolytic myofibers and reduced proportion of oxidative myofibers. Molecular analysis revealed that PRMT5 promotes methylation of mSREBP1a (mature SREBP1a) in skeletal muscles to upregulate lipogenic genes. Moreover, PRMT5 epigenetically represses the transcription of *Pnpla2* gene (encoding ATGL) via H4R3Me2s. Consistently, ablation of *Pnpla2* gene in *Prmt5^MKO^* mice (muscle-specific *Prmt5/Pnpla2* double KO) fully rescued the contractile phenotypes of the *Prmt5^MKO^* mice. Collectively, these data establish a novel function of PRMT5 as a regulator of lipid metabolism in myocytes that subsequently affect muscle development and contractile function.

## Results

### Myofiber-specific *Prmt5* KO reduces muscle mass in adult mice

We first examined *Prmt5* gene expression in postnatal hindlimb muscles. Among the 7 PRMTs, mRNA level of *Prmt5* was significantly higher than other isoforms (Supplementary Fig. S1A). In addition, *Prmt5* mRNA level in the skeletal muscle was the highest among various human tissues, though every tissue expressed *Prmt5* abundantly (Supplementary Fig. S1B)^39^. We also assessed the expression of *Prmt5* during muscle satellite cell differentiation in publicly available single-cell RNA sequencing (scRNA-seq) ^22^. Violin plots clearly highlighted that *Prmt5* expression is dramatically increased during differentiation correlated with *MyoG* expression (Supplementary Fig. S1C). We therefore hypothesized that PRMT5 plays a key role in the post-differentiation skeletal muscle. To test this hypothesis, we generated myocyte-specific *Prmt5* knockout mice (abbreviated as *Prmt5^MKO^*) by crossing *Prmt5^flox/flox^* mice with *Myl1^cre^* knockin mice expressing Cre recombinase driven by the endogenous *myosin light chain 1* (*Myl1*) gene. In this mouse model, the frameshift deletion of exon 7 causes a premature stop codon, resulting in a truncated protein of only 220 amino acids excluding all the key functional domain (Fig. 1A). A specific reduction of *Prmt5* in skeletal muscle was validated in *Prmt5^MKO^* mice in comparison to WT mice (Supplementary Fig. S1D).

**Figure 1:**
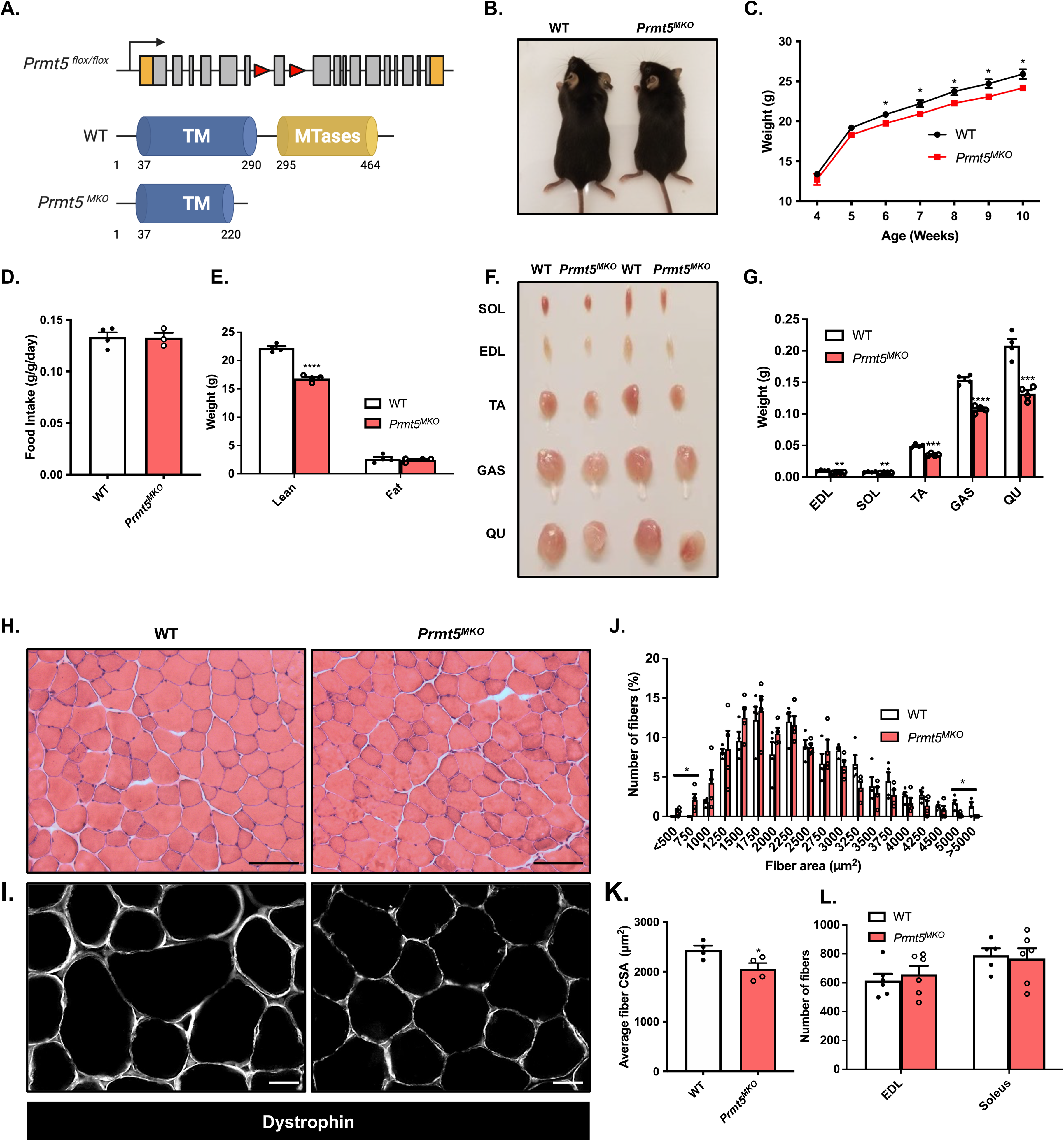
Muscle-specific knockout of PRMT5 (*Prmt5^MKO^*) leads to muscle mass reduction. **(A)** Genetic targeting strategy showing skeletal muscle-specific deletion of *Prmt5* using the *Cre-LoxP* recombinase under the control of the *Myl1* promoter. **(B,C)** Representative images of 2-month-old **(B)** and total body weight **(C)** of WT (n=6) *Prmt5^MKO^* mice (n=6) over time (right panel). (**D)**The measurement of food intake normalized to body weight in WT (n=4) and *Prmt5^MKO^* mice (n=3). **(E)** The measurement of lean and fat mass in WT (n=4) and *Prmt5^MKO^* mice (n=4) determined by using a EcoMRI body composition analyzer. **(F)** Photographs of skeletal muscles in 2-month-old WT and *Prm5*^MKO^ mice. **(G)**The quantified weight of muscles (SOL, EDL, TA, GAS, QU) from WT (n=4) and *Prmt5^MK¤^* mice (n=4). **(H,I)** A representative H&E **(H)**and immunofluorescence **(I)** of dystrophin staining in TA muscle cross-sections in WT and *Prmt5^MKO^* mice. Scale bar: 100 μm. **(J,K)** Distribution of myofiber size **(J)**and average cross-sectional area (CSA) of myofibers size **(K)**in TA muscle from WT (n=4) and *Prmt5^MKO^* mice (n=4). **(L)** The total number of myofibers in EDL and Soleus from WT (n=5) and *Prmt5^MKO^* mice (n=6). The data are presented as mean ± S.E.M in **C-E,G,J** and **K.** The *p* values determined by two-tailed ANOVA unpaired *t* test are indicated in **C,E,G,J** and **K**. The total number of biologically independent samples are indicated in **C-E,G,J, K** and **L**.

The *Prmt5^MKO^* mice were born normally but appeared leaner than their WT littermates starting at 6-week-old, despite similar food intake (Fig. 1B-D). EcoMRI body composition measurements revealed that the reduced body weights in the 2-Month-old KO mice is associated with significant reduction in lean mass (muscle) but not in fat mass (Fig. 1E). Consistently, the 2-month-old *Prmt5^MKO^* mice had smaller skeletal muscles, including Sol (Soleus), EDL (Extensor digitorum longus), TA (Tibialis anterior), GAS (Gastrocnemius), and QU (Quadriceps) muscles, than WT counterparts (Fig. 1F, G). In contrast, fat masses of various depots including iWAT (Inguinal white adipose tissue) and BAT (Brown adipose tissue) were similar between the WT and KO mice (Supplemental Fig. S1E). We also measured myofiber size based on cross-sectional area (CSA) in TA muscle sections after H&E staining and immunofluorescence staining of dystrophin (Fig. 1H, I). A leftward shift in the distribution of myofiber sizes was observed in 2-month-old *Prmt5^MKO^* mice (Fig. 1J), and the average CSA of the TA muscle in *Prmt5^MKO^* mice was significantly smaller than that of WT muscles (Fig. 1K). However, the total number of myofibers was identical between the WT and KO muscles (Fig. 1L). The observation that *Prmt5* KO only reduces myofiber size without affecting myofiber numbers suggest the *Prmt5^MKO^* mice does not affect myogenesis and only affect postnatal growth and maintenance of myofibers.

### *Prmt5* KO impairs motor performance and muscle contractile function

We examined muscle functions based on several physiological measurements. The grip strength of *Prmt5^MKO^* mice was significantly weaker than that of WT mice (Fig. 2A). When mice were run on a treadmill, the *Prmt5^MKO^* mice exhibited significantly lower maximum speed, running time, and running distance than did their littermate controls (Fig. 2B-D). To explore whether the reduced grip strength and exercise performance of *Prmt5^MKO^* mice was due to attenuated muscle contractile function, we assessed force generation capacity of the isolated fast-twitch EDL and slow-twitch Soleus muscles. Force development of the EDL and Soleus muscles was measured over a range of stimulation frequencies (Fig. 2E, F). The maximum absolute force of the EDL and Soleus muscles was reduced by 40% and 22%, respectively, in the KO mice relative to WT animals (Fig. 2E, F). A similar trend was observed in specific force, when absolute force was normalized to muscle CSA (Fig. 2G, H). The maximal specific forces of *Prmt5^MKO^* EDL and Soleus were 27% and 16% lower than the corresponding WT muscles, respectively (Fig. 2G, H). These observations indicate that PRMT5 loss-of-function leads to skeletal muscle weakness.

**Figure 2:**
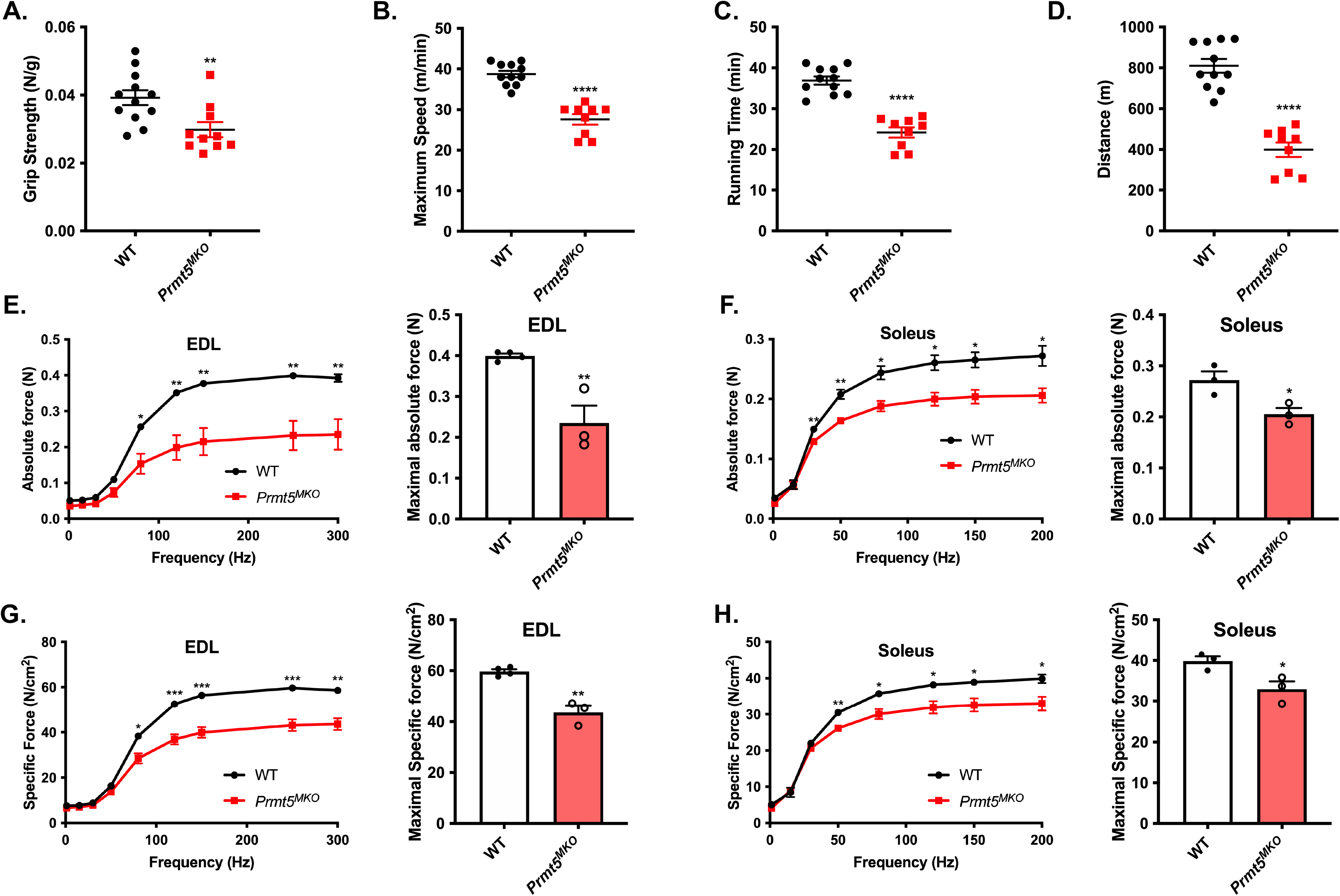
Reduced motor-performance and muscle contractile function in *Prmt5^MKO^* mice. **(A)** Grip strength tests of WT and *Prmt5^MKO^* mice assessed by grip force normalized to body weight WT (n=12) and *Prmt5^MKO^* mice (n=10). (**B,C,D)** Exercise performance of maximum speed **(B)**, running time **(C)**, running distance **(D)** of WT (n=11) and *Prmt5^MKO^* mice (n=9) measured by treadmill. **(E,F)** A graph of absolute **(E)** and specific force **(F)** (on the left panel) and maximal absolute and specific force (on right panel) on EDL muscle from WT (n=4) and *Prmt5^MKO^* mice (n=3). **(G,H)** A graph of absolute **(G)** and specific force **(H)** (on the left panel) and maximal absolute and specific force (on right panel) on Soleus muscle from WT (n=4) and *Prmt5^MKO^* mice (n=3). The data are presented as mean ± S.E.M in **A-H.** The *p* values by two-tailed ANOVA unpaired *t* test are indicated in **A-H**. The total number of biologically independent samples are indicated in **A-H**.

### *Prmt5* KO reduces oxidative myofibers while increasing glycolytic myofibers

Metabolic properties of myofibers have profound effects on exercise endurance and systemic metabolism^40^. We next assessed myofiber composition of representative fast (EDL) and slow (Soleus) muscles based on immunofluorescent staining of myosin heavy chain isoforms (Fig. 3A). A reduced proportion of oxidative type IIA myofibers and an increased proportion of glycolytic type IIB myofibers were observed in EDL and Soleus muscles of *Prmt5^MKO^* compared to WT muscles (Fig. 3B, C). We further carried out succinate dehydrogenase (SDH) staining as an indicator of mitochondrial oxidative capacity (Fig. 3D). Compared tothe WT EDL muscles, *Prmt5^MKO^* muscles had a lower abundance of SDH^high^ and SDH^med^ myofibers and a higher abundance of SDH^low^myofibers (Fig. 3E). The *Prmt5^MKO^* Soleus muscles also had reduced abundance of SDH^med^ myofibers and increased abundance of SDH^low^ myofibers (Fig. 3F). To evaluate the consequence of myofiber type alteration on whole-body metabolism, we used indirect calorimetry to measure the oxygen consumption (VO2) and carbon dioxide production (VCO_2_) (Fig. 3G, H). *Prmt5^MKO^* mice showed lower levels of oxygen consumption (VO_2_), carbon dioxide production (VCO_2_) and the heat production than WT counterparts, during both day and night (Fig. 3G, H, Supplementary Fig. S2A). However, there was no significant differences in the VO_2_ and VCO_2_ between WT and *Prmt5^MKO^* mice when the values were normalized to lean masses (Supplementary Fig. S2B-C), due to the reduced lean masses in the KO mice (see Fig. 1E). Collectively, these findings suggest that *Prmt5^MKO^* alters muscle metabolism by shifting oxidative myofibers toward glycolytic myofibers.

**Figure 3:**
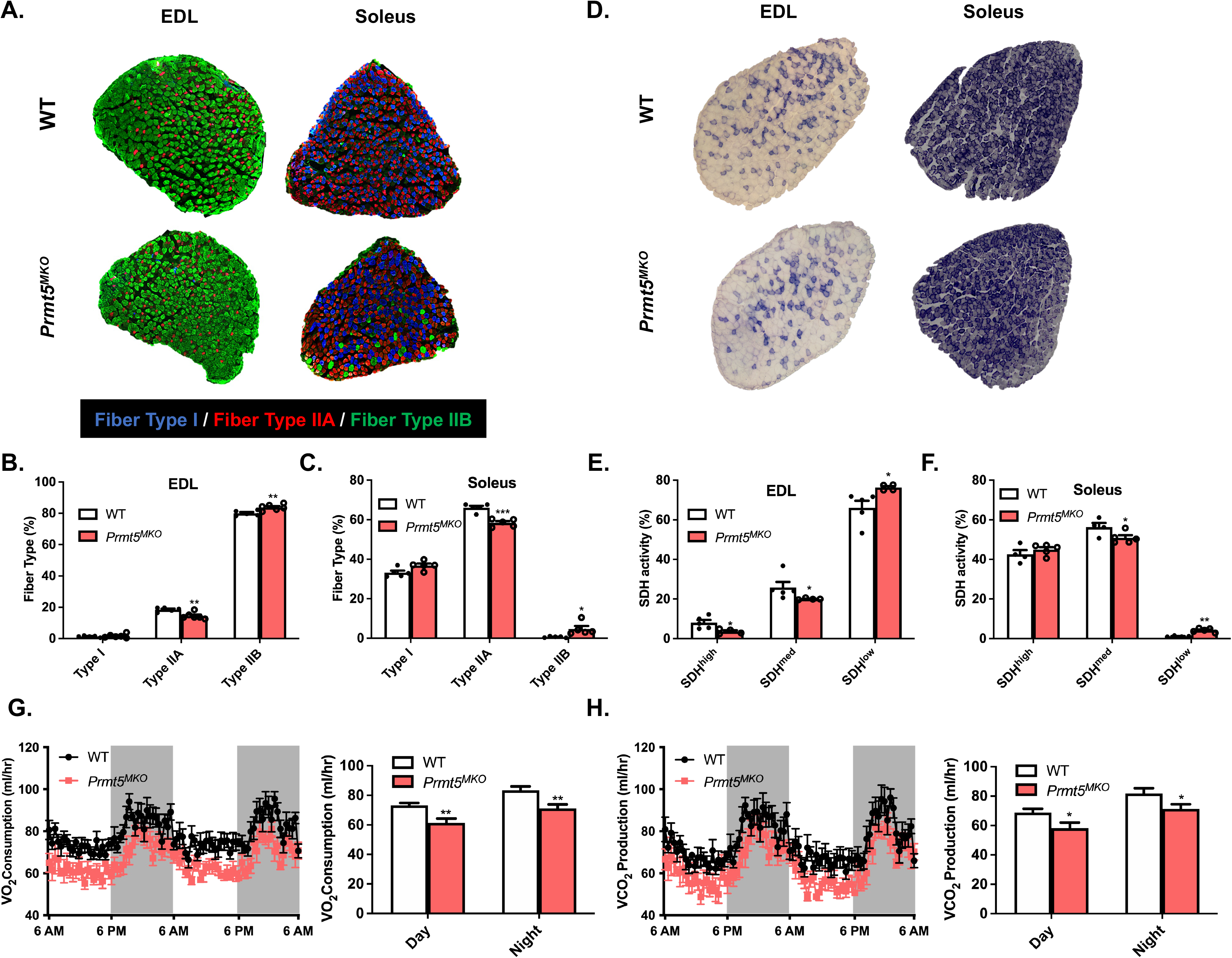
Depletion of *Prmt5* leads to a fiber-type switch toward glycolytic myofibers but does not affect systemic metabolism. **(A)** Representative immunostaining of 2-month-old WT and *Prmt5^MKO^* mice for 3 fiber types (fiber type I, fiber type IIA, fiber type IIB) in EDL and Soleus muscles. **(B,C)** Quantification of stained 3 fiber-type in EDL **(B)** and Soleus **(C)** muscles from WT and *Prmt5^MKO^* mice; (n=5). **(D)** Representative histochemical staining image of succinate dehydrogenase (SDH) enzymatic activity in EDL and Soleus muscles. **(E,F)** Quantification of stained myofibers by 3 different staining grades in EDL **(E)** and Soleus **(F)** muscles from WT (n=5) and *Prmt5^MKO^* mice (n=4). **(G,H)** Metabolic rate of O_2_ consumption **(G)** and CO_2_ production **(H)** measured by an indirect calorimetry of 4-6-month-old WT (n=8) and *Prmt5^MKO^* (n=7) mice. The data are presented as mean ± S.E.M in **B-C,** and **E-H.** The *p* values by two-tailed ANOVA unpaired *t* test are indicated in **B-C,** and **E-H**. The total number of biologically independent samples are indicated in **B-C**, and **E-H**.

### PRMT5 regulates lipid metabolism in skeletal muscles

Given the fiber type switching in *Prmt5^MKO^* mice and well-known metabolic differences among fiber types, we further explored lipid metabolism which is more pronounced in oxidative myofibers^41^. As the first step, we assessed neutral lipid content by Oil Red O (ORO) staining in TA muscle sections and observed in WT muscles under brightfield imaging clusters of myofibers with numerous ORO^+^ lipid droplets, presumably representing oxidative myofibers (Fig.4A). However, ORO+ labeled lipid droplets were completely absent in the *Prmt5* KO muscles (Fig. 4A). We also used more sensitive fluorescent imaging (as ORO emits red fluorescence) and again observed fluorescent puncta representing lipid droplets in selective clusters of WT myofibers (Fig. 4B). In contrast, ORO puncta were absent in *Prmt5^MKO^* muscles, and only weak and non-puncta ORO fluorescence were detectable (Fig. 4B). These data demonstrate that *Prmt5* KO deregulates lipid metabolism in myofibers.

**Figure 4.**
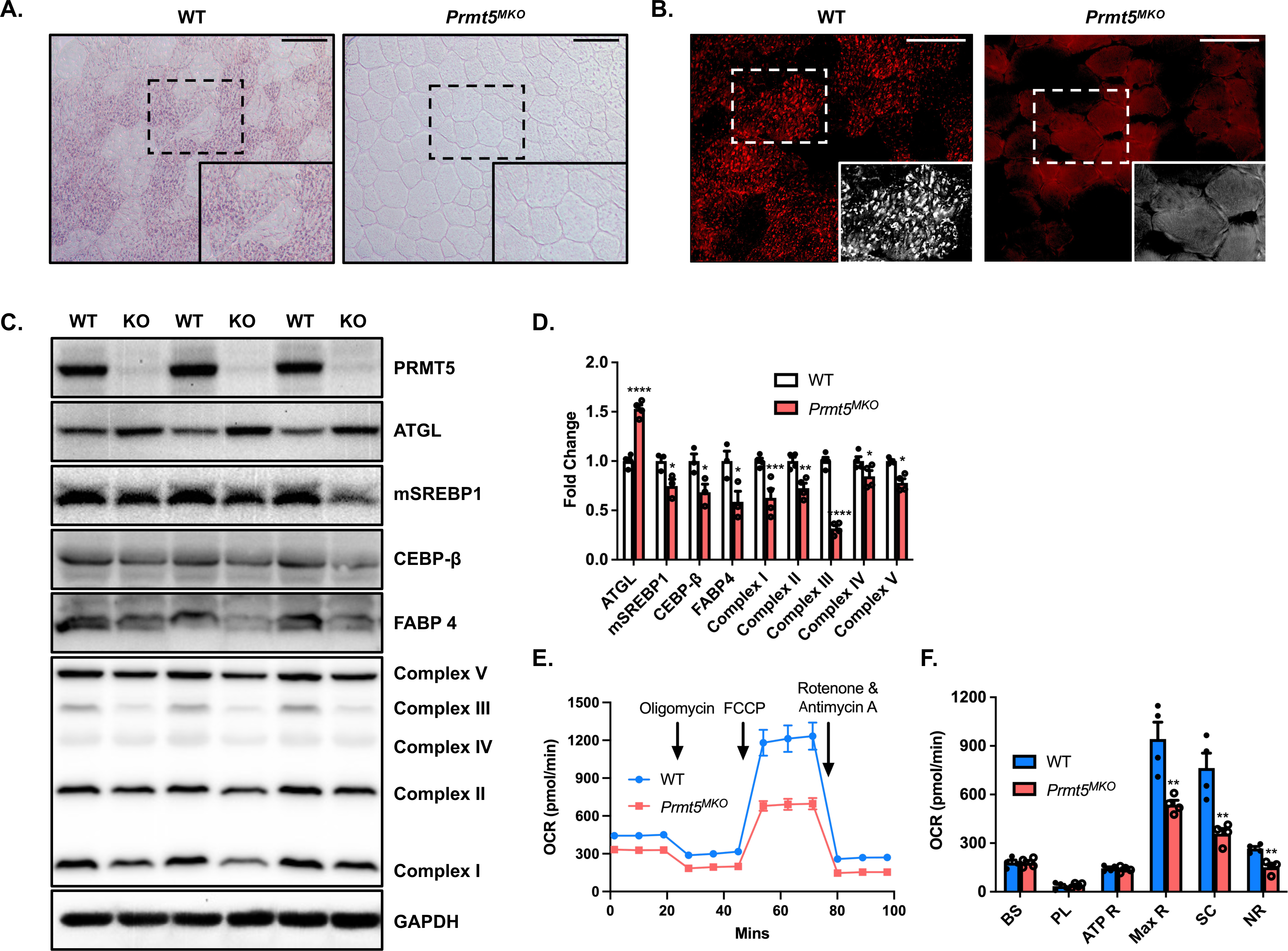
PRMT5-deficient skeletal muscles exhibit lower lipid content and metabolic rate of oxygen consumption rate (OCR) **(A,B)** Representative images of ORO staining **(A)** and immunofluorescent ORO staining **(B)** in TA muscle from WT and *Prmt5^MKO^* mice, Scale bar: 100 μm. **(C)** Western blotting analysis for protein markers of lipolysis, lipogenesis, and electron transport chain (ETC) complexes in skeletal muscle from WT (n=3) and *Prmt5^MKO^* mice; (n=3). **(D)** Quantification of relative protein levels of ATGL, mSREBP1, CEBP-β, FABP4, OXPHOS complexes (normalized to GAPDH) in skeletal muscles isolated from WT and *Prmt5^MKO^* mice; (n=3). **(E)** Seahorse horse analysis for the measurement of oxygen consumption rate (OCR) in myotubes isolated from hindlimb muscles of 4-week-old WT and *Prmt5^MKO^* mice; (n=4). **(F)** OCR was measured at basal state and after sequential addition of Oligomycin, FCCP, and Rotenone/Antimycin A to determine basal respiration (BS), proton leak (PL), ATP respiration (ATP R), maximal respiration (Max R), spare capacity (SC), and non-mitochondrial respiration (NR); (n=4). The data are presented as mean ± S.E.M in **D-F.** The *p* values by two-tailed ANOVA unpaired *t* test are indicated in **D-F.** The total number of biologically independent samples are indicated in **D-F**.

We further examined the levels of several key proteins involved in lipid metabolism (ATGL/*Pnpla2*, mSREBP1, CEBP-β, FABP4) and downstream mitochondrial electron transport chain (ETC) mediating FAO (Fig. 4C). Quantification results showed that the levels of ATGL, a rate-limiting enzyme of lipolysis, were consistently higher but the lipogenic and mitochondrial ETC proteins were all lower in *Prmt5^MKO^* compared to WT muscles (Fig. 4D). We investigated how PRMT5-dependent lipid metabolism affects mitochondrial respiration through Seahorse Analysis (Fig. 4E). The results showed that *Prmt5^MKO^* led to significantly lower levels of oxygen consumption associated with maximal respiration (Max R) and spare capacity (SC) (Fig. 4F). Overall, these findings demonstrate that PRMT5 is crucial for maintaining normal lipid metabolism and mitochondrial respiratory activity in the skeletal muscles.

### PRMT5 methylates mSREBP1a to increase its stability

To understand how PRMT5 regulates lipid metabolism in myofibers, we focused on mSREBP1a, a key transcriptional factor regulating lipogenesis. As *Prmt5* KO reduced the level of mSREBP1 (Fig. 4C, D), we hypothesized that PRMT5 stabilizes mSREBP1a. We expressed PRMT5-GFP and SREBP1a-Flag fusion proteins in C2C12 myoblasts and confirmed the overexpression of corresponding proteins (Fig. 5A). We then performed co-immunoprecipitation (co-IP) using Flag antibody and observed that PRMT5 not only binds to mSREBP1a, but methylates mSREBP1a (Fig. 5A). Conversely, co-IP using GFP antibody (for PRMT5-GFP) also confirmed pull-down of mSREBP1-Flag (Fig. 5B). We also performed co-IP and western blots (WB) on WT and *Prmt5^MKO^* muscles lysates, showing reduced levels of total mSREBP1 and symmetrically dimethylated mSREBP1 in the KO samples (Fig. 5C). Notably, the ratio of symmetrically dimethylated mSREBP1 to total mSREBP1 was significantly lower in the *Prmt5^MKO^* than in the WT muscles (Fig. 5D).

**Figure 5.**
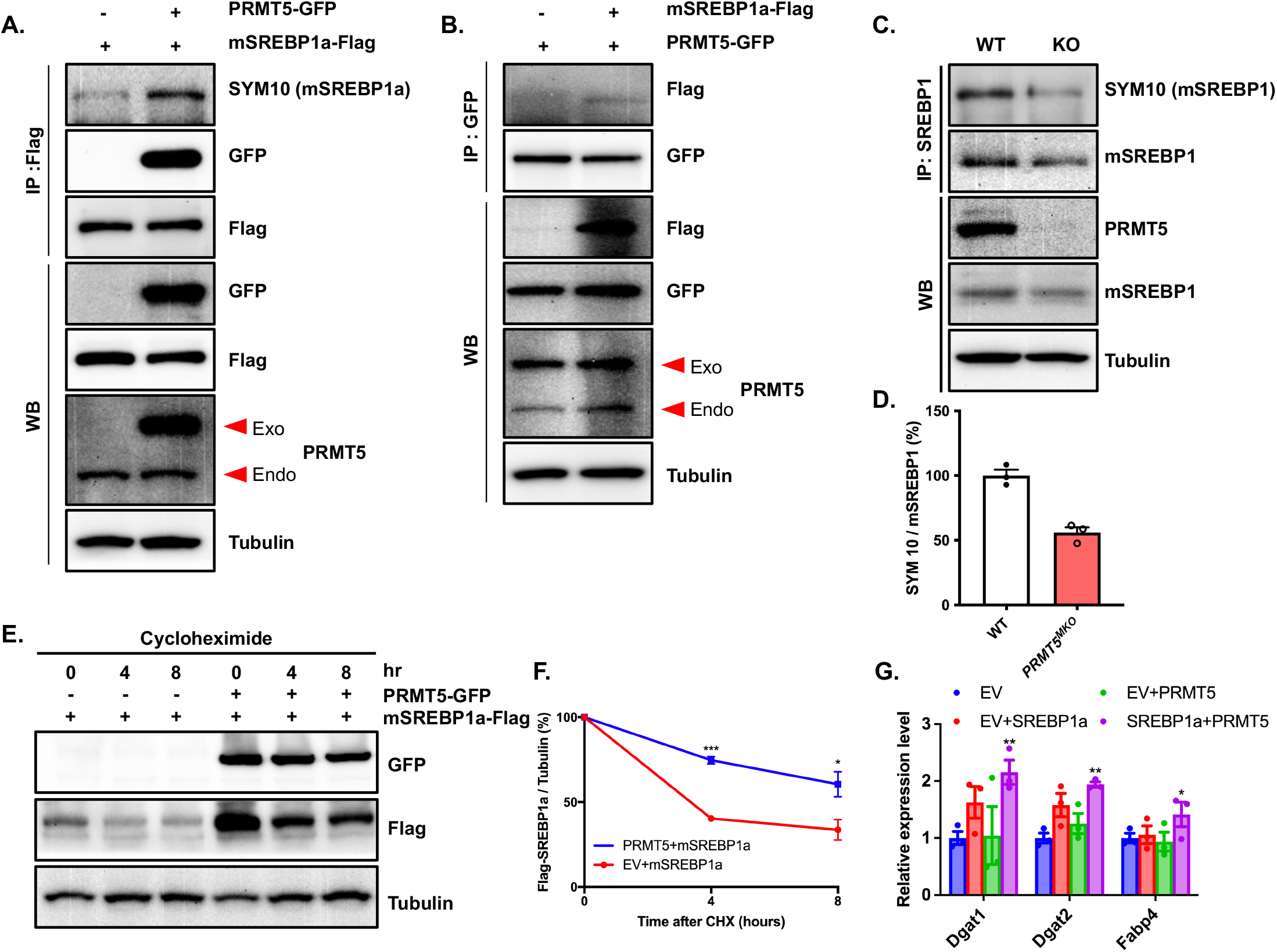
PRMT5 stabilizes regulates lipid accumulation through ATGL and SREBP1a methylation. **(A)** C2C12 cells overexpressing PRMT5-GFP or PRMT5-GFP and SREBP1a-Flag were immunoprecipitated with Flag antibody and blotted with SYM10, GFP, Flag, PRMT5, and Tubulin antibodies. **(B)** C2C12 cells overexpressing SREBP1a-Flag or SREBP1a-Flag and PRMT5-GFP were immunoprecipitated with GFP antibody and blotted with Flag, GFP, PRMT5, and Tubulin antibodies. **(C)** Protein extracts of skeletal muscle isolated from WT and *Prmt5^MKO^* mice were immunoprecipitated with SREBP1 antibody and immunoblotted with SYM10 (mSREBP1), SREBP1, PRMT5, and tubulin antibodies. **(D)** Quantification of relative methylated mSREBP1 (normalized to mSREBP1) in **(C); (** n=3). **(E)** HEK293 cells were transfected with PRMT5-GFP or PRMT5-GFP and SREBP1a-Flag for 24 hours, followed by 0,4,8 hours cycloheximide (30 μg/ml). Lysates were immunoblotted with Flag, GFP, and tubulin antibodies. **(F)** Intensity of Flag was normalized to tubulin, then normalized to 0 hr; (n=3). **(G)** Relative expression of lipogenesis genes (*Dgat1, Dgat2, Fabp4*) in C2C12 transfected with PRMT5 and SREBP1a; (n=3). Data are presented as mean ± S.E.M in **D,F** and **G.** The *p* values determined by two-tailed ANOVA unpaired *t* test are indicated in **D,F** and **G.** The total number of biologically independent samples are indicated in **D,F** and **G**.

To explore the biological significance of PRMT5 mediated dimethylation of mSREBP1a, we performed protein stability assay. We overexpressed mSREBP1a with or without PRMT5 in C2C12 myoblasts (Fig. 5E). Cells were then treated with cycloheximide (CHX) to inhibit new protein synthesis. Proteins were collected at 0, 4, 8 h after addition of CHX to determine their degradation (Fig. 5E). Analysis of mSREBP1a levels over time indicates that PRMT5 expression significantly increased the stability of SREBP1a (Fig. 5F). We also analyzed the expression of genes responsible for TAG synthesis and fatty acid transport (*Dgat1, Dgat2, Fabp4*) in the C2C12 cells overexpressing mSREBP1a along or together with PRMT5 (Fig. 5G). The results show that PRMT5/SREBP1a co-expression significantly elevated the levels of *Dgat1, Dgat2*and *Fabp4* compared to SREBP1a overexpressing group (Fig. 5G). These results together demonstrate that PRMT5-mediated dimethylation stabilizes mSREBP1a to increase its transcriptional activity.

### PRMT5 mediates repressive H4R3 dimethylation to regulate *Pnpla2* expression

We next explored how PRMT5 regulates ATGL (encoded by *Pnp/a2* gene) expression and lipolysis in muscle cells. PRMT5 has been reported to symmetrically dimethylates arginine residues in histones to repress gene transcription^30–32^. We found that symmetric dimethylation of H4R3 (H4R3Me2s) was significantly reduced in *Prmt5^MKO^* muscle tissues (Fig. 6A, B). In addition, *Pnpla2* expression was upregulated in *Prmt5^MKO^*muscle tissues but down-regulated in PRMT5-overexpressing C2C12 myoblasts (Fig. 6C, D). These data prompted us to hypothesize that PRMT5 directly regulates transcription of *Pnpla2* through repressive H4R3Me2s. To test this, we analyzed a ChIP-seq dataset^42^ and identified a PRMT5 binding peak at around 3,000 bp upstream of transcription start site (TSS) of the *Pnp/a2* gene (Fig. 6E). ChIP-qPCR analysis confirmed that this region is highly enriched by both PRMT5 and H4R3Me2s in newly differentiated C2C12 myotubes (Fig. 6F, G). To substantiate this finding, another ChIP-qPCR assay was performed using WT and *Prmt5^MKO^* muscle tissues, and *Prmt5^MKO^* diminished the binding of PRMT5 and H4R3Me2s at the *Pnpla2* promoter region (Fig. 6H, I). Taken together, these results demonstrate that *Pnpla2* is epigenetically repressed by PRMT5 through H4R3Me2s.

**Figure 6:**
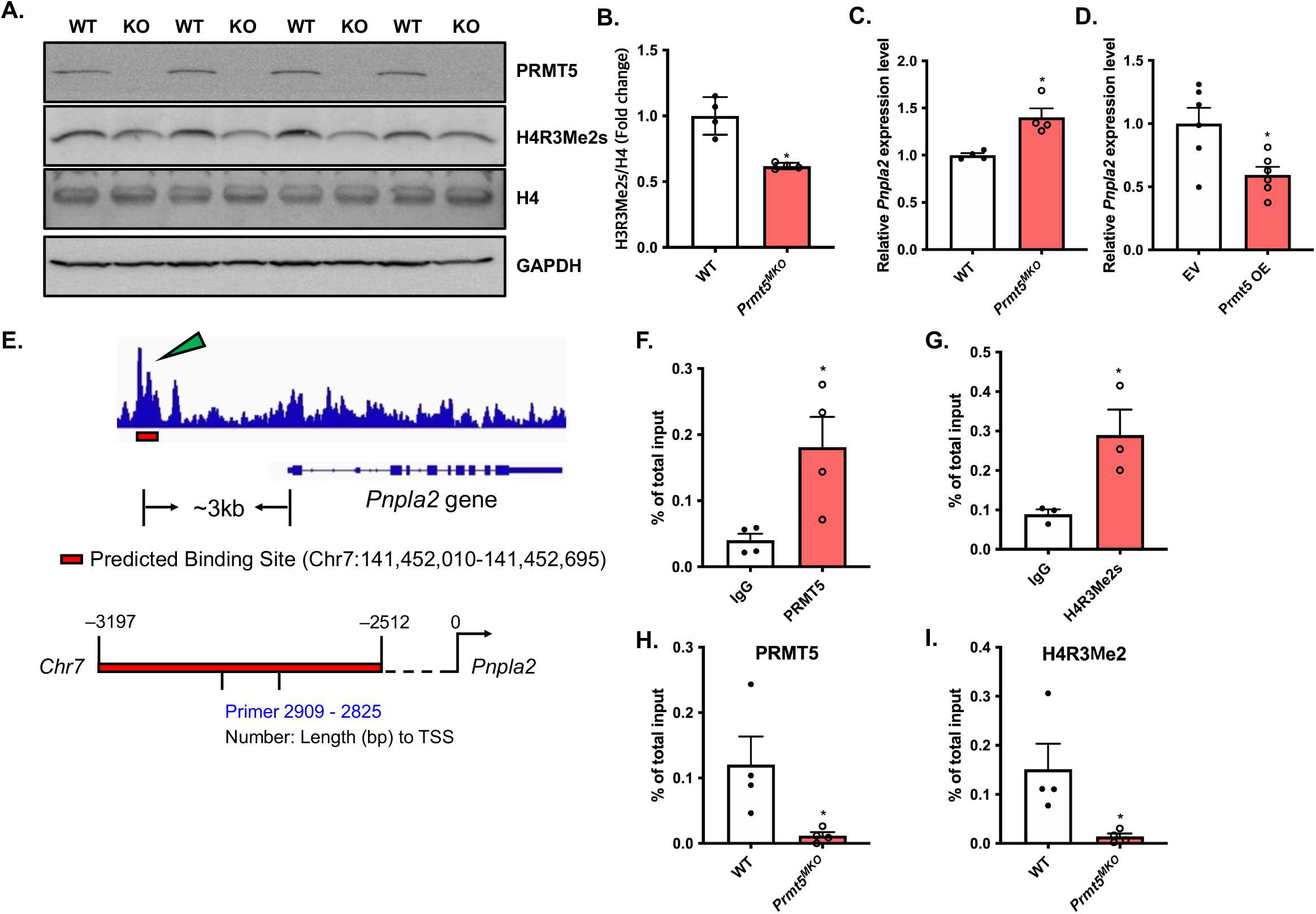
Epigenetic repression of *Pnpla2* transcription by PRMT5 in skeletal muscle. **(A,B)** Immunoblots showing dimethylation of H4R3 in skeletal muscle from WT (n=4) and *Prmt5^MKO^* mice (n=4) **(A)**, and quantification of H4R3Me2s normalized to total H4 **(B)** Relative *Pnpla2* expression in skeletal muscle from WT and *Prmt5^MKO^* mice; (n=4) **(C)** Relative *Pnpla2* expression in PRMT5-overexpressed C2C12 myoblasts; (n=6). **(E)** Chip-sequencing results showing the *Prmt5* binding peak on the *Pnpla2* promoter. **(F,G)** Enrichment of PRMT5 **(F)** and H4R3Me2s **(G)** on the proximal promoter region of the *Pnpla2* gene in myotubes; (n=4 for **F**, n=3 for **G**). **(H,I)** Enrichment of PRMT5 **(H)** and H4R3Me2s **(I)** on the proximal promoter region of the *Pnpla2* gene in skeletal muscle isolated from WT (n=4) and *Prmt5^MKO^* mice (n=4). The data are presented as mean ± S.E.M in **B-D,** and **F-I.** The *p* values by two-tailed ANOVA unpaired *t* test are indicated in **B-D,** and **F-I.** The total number of biologically independent samples are indicated in **B-D,** and **F-I**.

### *Pnpla2* KO normalizes muscle mass and function of the *Prmt5^MKO^* mice

ATGL upregulation drives muscle wasting in cancer cachexia^18^. To directly test if upregulation of ATGL is responsible for the phenotypes of the *Prmt5^MKO^* mice, we used *Myl1^cre^* to drive muscle-specific double KO of *Prmt5* and *Pnpla2* (*Prmt5/Pnpla2^MKO^*),Western blotting confirmed the loss of PRMT5 and ATGL in *Prmt5/Pnpla2^MKO^* muscles (Supplementary Fig. S3A). The loss of *Pnpla2* restored the levels of mitochondrial ETC proteins in *Prmt5* KO tissue (Supplementary Fig. S3A). Strikingly, the *Prmt5/Pnp/a2^MKO^* mice appeared similar to WT mice, both are larger than the *Prmt5^MKO^* mice (Fig. 6A). The body weights and lean masses of the *Prmt5/Pnpla2^MKO^* mice were also comparable to those of the WT mice, and higher than those of the *Prmt5^MKO^* mice (Fig. 6B, C). The myofiber size was also restored to the WT level in the *Prmt5/Pnpla2^MKO^* mice (Supplementary Fig. S3B-D). Performance on the incremental treadmill test and limb grip strength in *Prmt5/Pnpla2^MKO^* mice were also restored to the WT level (Fig. 6D, E). Moreover, the absolute and specific forces of the EDL and Soleus muscles of *Prmt5/Pnpla2^MKO^* were significantly higher than that of the *Prmt5^MKO^* mice, with the maximal force of *Prmt5/Pnplal2^MKO^* mice muscles being identical to that of WT mice (Fig. 6F-I). We also assessed lipid content by ORO staining in myofibers and found a clear increase in the number of ORO^+^ LDs in *Prmt5/Pnpla2^MKO^* muscles compared to *Prmt5^MKO^* muscles under brightfield and fluorescent imaging (Supplementary Fig. S3E). Collectively, these data demonstrate that PRMT5 regulates neutral lipid content, myocyte size and function through ATGL.

## Discussion

Our study uncovers a new role of PRMT5 in myofiber metabolism and contractile function. Utilizing a muscle-specific *Prmt5* KO mouse model, we provided physiological, histological, and molecular evidence to support that PRMT5 ablation leads to muscle atrophy and favors the formation of glycolytic myofibers. Mechanistically, PRMT5 on one hand mediates mSREBP1a methylation to stabilize this protein and promote lipogenesis, on the other hand mediates H4R3me2 to repress *Pnpla2* expression in myofibers, thus limiting lipolysis. Thus, PRMT5 should normally promote IMCL deposition through stimulating lipogenesis and limiting lipolysis. Subsequently, concomitant deletion of *Pnpla2* restored muscle functions of the *Prmt5^MKO^* mice (Fig. 8). These results establish a new role of PRMT5 in linking the contractile and metabolic function of myocytes.

**Figure 7:**
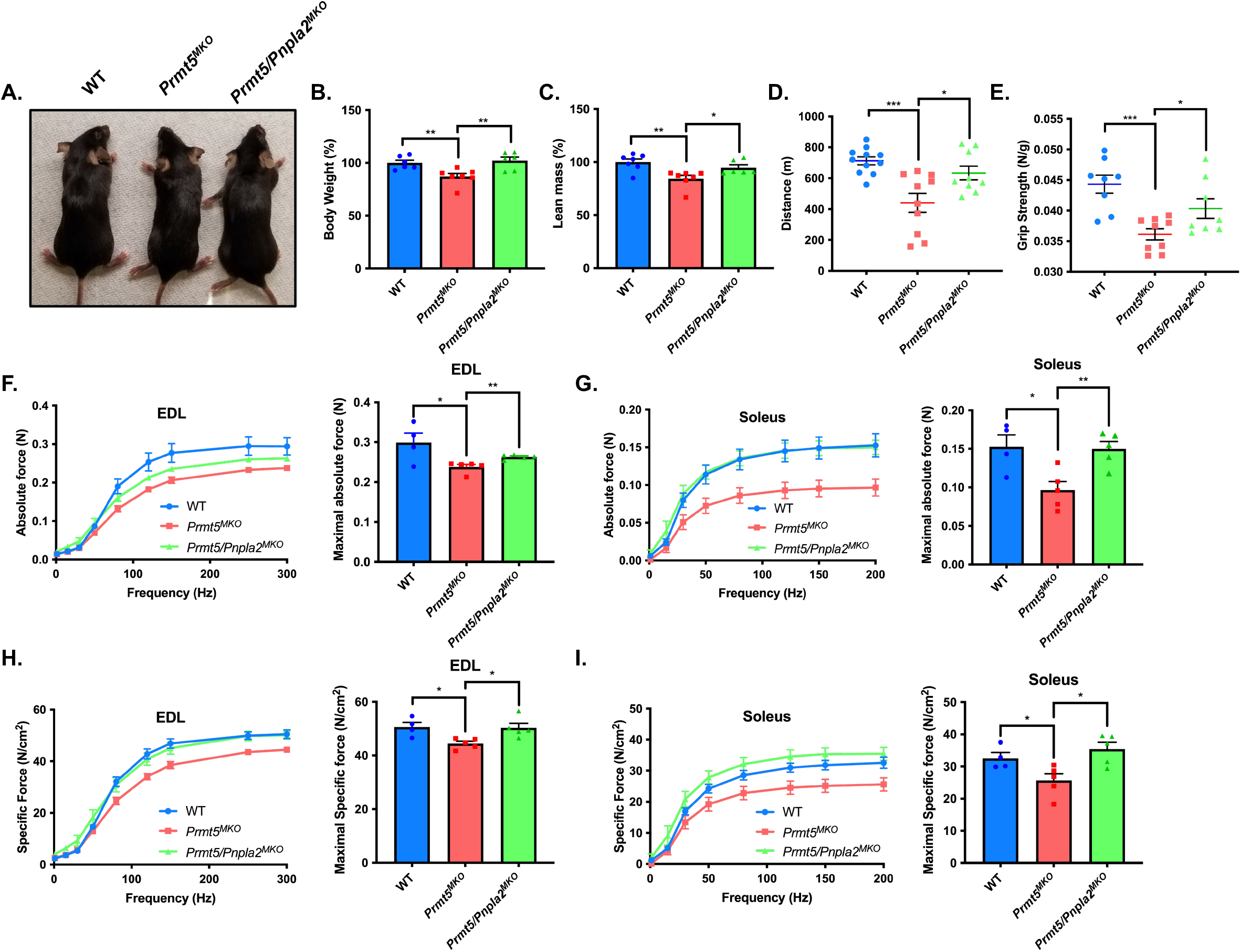
*Pnpla2* deletion restores muscle function of *Prmt5^MKO^* mice. **(A,B)** A representative images of whole body **(A)** and the measurement of body weight **(B)** of WT (n=6), *Prmt5^MKO^* mice (n=7) and *Prmt5/Pnpla2^MKO^* mice (n=6) at 8 weeks of age. **(C)** Percentage of lean mass determined by body composition analyzer for WT (n=6), *Prmt5^MKO^* mice (n=7) and *Prmt5/Pnpla2^MKO^* mice (n=6) at 8-weeks-old. **(D,E)** The assessment of exercise performance in running distance **(D)** and griping strength test **(E)** in WT (n=11 for **D**, n=8 for **E**), *Prmt5^MKO^* mice (n=10 for **D**, n=9 for **E**) and *Prmt5/Pnpla2^MKO^* mice (n=9 for **D**, n=8 for **E). (F,G)** A graph of absolute force **(left panel)** and maximal absolute force **(right panel)** of EDL **(F)** and Soleus **(G)** muscles from WT (n=4), *Prmt5^MKO^* mice (n=5) and *Prmt5/Pnpla2^MKO^* mice (n=5). **(H,I)** A graph of specific force **(left panel)** and maximal specific force **(right panel)** of EDL **(H)** and Soleus **(I)** muscles from WT (n=4), *Prmt5^MKO^* mice (n=5) and *Prmt5/Pnpla2^MKO^* mice (n=5). The data are presented as mean ± S.E.M in **B-I**. The *p* values by two-tailed ANOVA unpaired *t* test are indicated in **B-I** . The total number of biologically independent samples are indicated in **B-I**.

**Figure 8:**
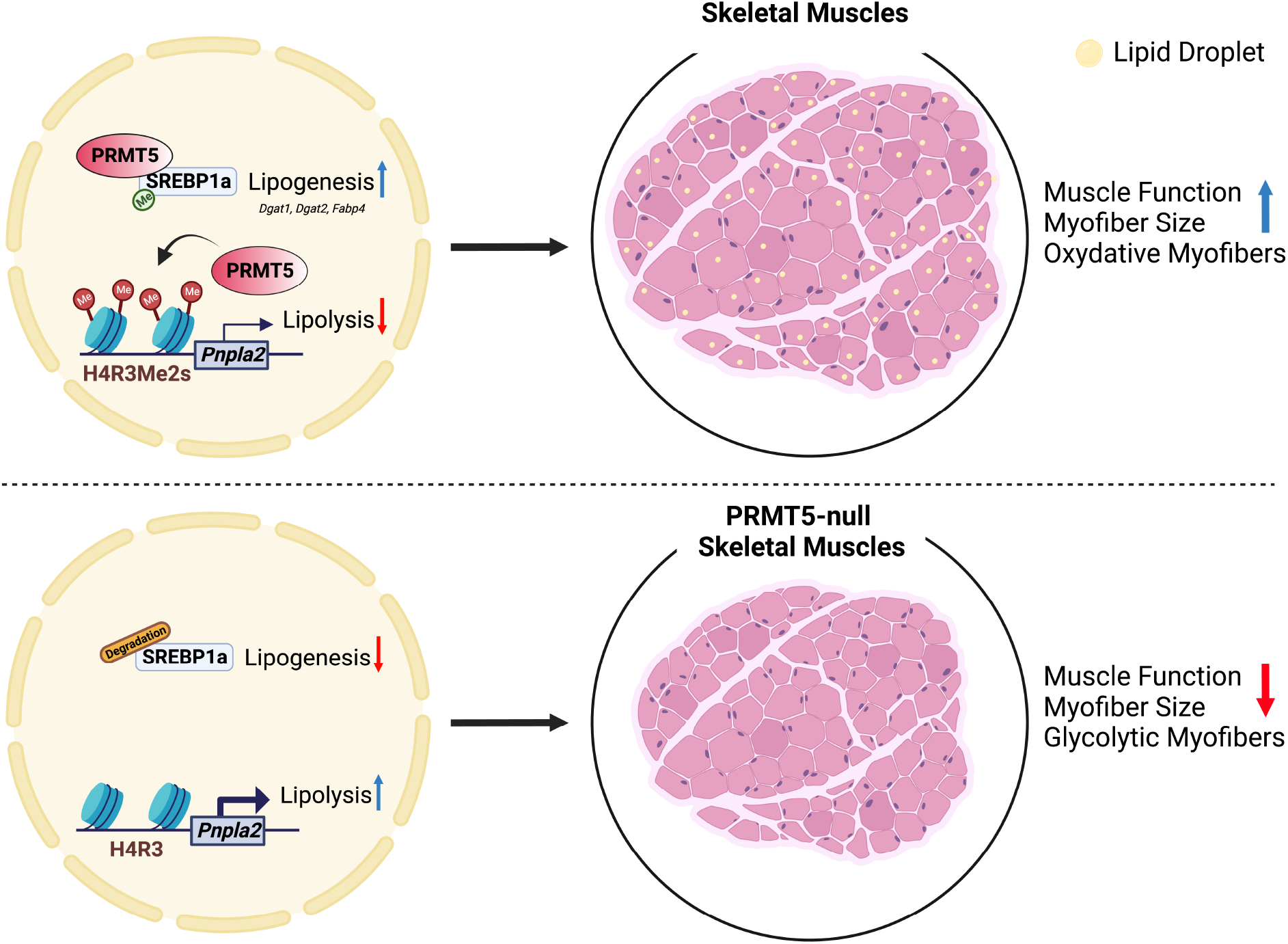
A schematic model depicting the role of PRMT5 in muscle function. PRMT5 is highly expressed in skeletal muscle and necessary to mediate symmetric dimethylation of H4R3 at the *Pnpla2* gene and to stabilize mSREBP1a to promote LD formation. Gene inactivation of *Prmt5 in vivo* attenuates physical activity and decreases myofiber size accompanied by reduced LD deposition. The deletion of *Pnpla2 in vivo* can restore the decreased muscle function by *Prmt5* loss.

Previous work has reported a key role of PRMT5 in regulating proliferation of muscle progenitor cells, and the deletion of *Prmt5* impairs progenitor cell-mediated muscle regeneration^38^. *In vitro* studies on PRMT5 have revealed its role in regulating COPR5-mediated cell proliferation and BRG1/MyoD-dependent chromatin remodeling for myogenic activation within myoblasts^37,43^. Although PRMT5 level has been reported to be influenced by exercise in rodent and human skeletal muscles^44,45^, the specific role of PRMT5 in myofibers has been unknown. This represents a critical knowledge gap given the elevated *Prmt5* expression during myogenic differentiation (supplemental Fig. S1). In this work, we used muscle specific *Myl1^cre^* mice to drive the deletion of *Prmt5* in postdifferentiation myocytes. *Myl1* gene is activated at E9 during embryonic development within most skeletal muscles *in vivo* and is expressed in mononuclear myocytes during differentiation *in vitro*^46,47^, Therefore, the muscle atrophy of the *Prmt5^MKO^* mice is independent of myogenic differentiation.

In our study, we observed the muscle atrophy of *Prmt5^MKO^* mice is associated with elevated levels of ATGL, which promotes lipolysis of lipid droplets in myocytes. PRMT5-dependent mSREBP1a methylation in nucleus is reported to upregulate *de novo* lipogenesis in adipocytes and cancer cells^42,48^. Our study clearly demonstrated that mSREBP1a is stabilized through methylation by PRMT5, and co-overexpression of *Prmt5* and *Srebp1a* upregulates lipogenic genes. Our results suggest that a normal lipid droplet content is necessary for proper function of the muscle. Lipid accumulation in myocytes has been associated with functional declines of muscle in lipid storage myopathies^49^. Our results demonstrate that a lack of lipids in the myocytes severely compromises muscle contractile function and reduces muscle mass. Our study further suggests that targeting PRMT5 using commercially available pharmacological inhibitors could be a potential therapy for patients with lipid myopathies.

Muscle fiber type switching occurs in response to a variety of genetic, physiological, and metabolic condition^50^. A decrease in the proportion of slow-oxidative myofibers is, in general, correlated with reduced exercise plasticity^51^. Several PRMTs have been reported to affect fiber type composition^52^. An increase in the proportion of glycolytic fibers was reported in *Prmt7* KO mice, resulting in reduced endurance exercise capacities^35^. In our study, loss of PRMT5 protein also converts oxidative myofibers to glycolytic fiber type, suggesting that PRMT5 is essential for specification or maintenance of the oxidative myofiber types and exercise endurance. Oxidative myofibers normally have a higher mitochondrial content than glycolytic myofibers to enable cellular energy production through oxidative phosphorylation^53^. In pharmacological and genetic inactivation studies *in vivo,* mitochondrial enzyme activity has been shown to be positively associated with exercise capacity^51,54^. Our finding indicated a reduction in oxidative phosphorylation in skeletal muscle tissues, as well as reduced OCR in *Prmt5^MKO^* myocytes. These metabolic alterations may have accounted for the impaired exercise capacity, as *Pnpla2* KO in *Prmt5^MKO^* mice rescues not only the lipid droplet content but also muscle contractile functions.

Gene expression and transcription initiation are regulated by numerous epigenetic markers^24,26^. An epigenetic role of PRMT5 has been shown to regulate chromatin remodeling through histone modification during myogenesis^43,55,56^. We show that in terminally differentiated myocytes, PRMT5-mediated H4R3Me2s represents a significant epigenetic regulatory mechanism underlying muscle function and metabolism. We reveal that PRMT5 represses *Pnpla2* expression through directly binding to its promoter. Consistent with our results, PRMT7 and PRMT5 mediates H4R3Me2 represses expression of *Dnmt3b and Flip1L*, respectively^57,58^. Interestingly, SIRT7-mediated desuccinylation at K387 enhances methyltransferase activity of PRMT5 by promoting formation of PRMT5-Mep50 complex to regulate lipid metabolism, and PRMT5 facilitates fatty acid biogenesis through mSREBP1a methylation and BSCL2-mediated lipid droplet formation through methylating SPT5 in adipocytes^42,59^. However, the protein levels of ATGL are positively regulated by PRMT5 in adipose tissue but negatively regulated by PRMT5 in myocytes, suggesting tissue-dependent roles of PRMT5^42^.

Although we show that PRMT5 represses lipolysis in myocytes, whether it also regulates mitochondrial fatty acid oxidation (FAO) remains unknown. Lipid dynamics are modulated by mitochondrial activity, and oxidation of FAs is linked to ATP production for proper muscle function^60^. Patients with lipid storage disorder show disrupted mitochondrial activity in respiratory complexes, and it leads to muscle wasting^16,61^. Furthermore, mitochondrial-targeted antioxidants improved muscle atrophy by preventing reactive oxygen (ROS) production^62^. Future studies will be needed to address the involvement of PRMT5 in regulating mitochondrial activity in myocytes.

## MATERIALS AND METHODS

### Mice

All procedures involving mice were performed in compliance with the institutional guidelines of Purdue University Animal Care and Use Committee. *Myl1^cre^* (Stock # 024713) mouse strains were provided by Steven Burden (Scribal Institute of Biomolecular Medicine, NYU), *Rosa26-Cre^ER^* (Stock # 008463) mouse strains were bought from Jackson Laboratory, and *Prmt5^flox/flox^* (Stock # 034414) and *Pnpla2*^flox/flox^ (Stock # 024278) mouse strains were obtained as described previously^22,42^. The genotypes of experimental WT and KO animals are as follows: WT (*Prmt5^flox/flox^*), *Prmt5^MKO^* (*Myl1^cre^; Prmt5^flox/flox^*) and *Prmt5/Pnpla2*^MKO^(*Myl1^cre^; Prmt5^flox/flox^; Pnpla2^flox/flox^*), The primers for genotyping are listed in Supplementary Table 1. Mice were housed in the animal facility with free access to water and standard rodent chow food or high fat diet (HFD, TD.06414 Harlan). 2-month-old mice were used unless otherwise indicated. Food intake was calculated by measuring weekly food consumption normalized to body weight in each cage

### Indirect calorimetry study

Oxygen consumption (VO_2_) and carbon dioxide production (VCO_2_) levels were assessed by using an indirect calorimetry system (Oxymas, Columbus instruments), as previously described^63^. Mice were individually housed in chambers and had free access to food and water under a constant environmental room temperature.

### Isolation and culture of primary myoblasts

Primary myoblasts were isolated from hind limb skeletal muscles of 4-week-old mice as previously described^64^. Muscle tissues were minced and digested in type II collagenase and dispase B mixture (Roche). Digestion was neutralized by adding growth media containing F-10 Ham’s medium (ThermoFisher Scientific), 20% fetal bovine serum (FBS), 1% penicillin, 4 ng/ml basic fibroblast growth factor (ThermoFisher Scientific), and cells were cultured on collagen-coated plates. Pre-plating was performed to purify primary myoblasts. For differentiation, primary myoblasts were plated on the BD Matrigel-coated culture plates and differentiated in DMEM supplemented with 2% horse serum and 1% penicillin.

### Cell cultures and transfection

C2C12 cells were cultured in Dulbecco’s modified Eagle’s medium (DMEM) supplemented with 10 % FBS (Hyclone) and 1 % penicillin, in a 37 °C humidified incubator with 5 % CO_2_. For differentiation, C2C12 cells were plated on the BD Matrigel-coated culture plates and differentiated in DMEM supplemented with 2% horse serum, 400 nM insulin, and 1% penicillin. Plasmids and lipofectamine 2000 mixture were transfected in C2C12, based on the manufacturer’s instructions, and harvested after 24 hours. For transfection, pcDNA-GFP-PRMT5 plasmid and pcDNA3.1-2xFLAG-SREBP1a plasmid were mixed with lipofectamine 2000 Reagent in Opti-MEM media (Giobco) and transfected into 60 % confluent cells following the protocol from manufacture. Opti0MEM media was changed to growth medium after 4 hours and cells were harvested after 24 hours for further analysis.

### scRNA-Seq data analysis

scRNA-seq datasets for muscle satellite cell differentiation were downloaded from publicly available data (GSE150366). For data analysis, barcodes and reads were aligned to mm10 (*Mus Musculus*) using CellRanger v3.1 and data analysis was performed using Seurat v3.1 as previously described^22^.

### H&E, immunofluorescence, and Oil red O (ORO) staining

Whole muscle tissues were immediately frozen in optical cutting temperature compound (OCT compound), and subsequently cut into 10 μm thick cross sections using a Leica CM1850 cryostat for H&E, immunofluorescence, and Oil red O staining. For H&E staining, the slides were stained in hematoxylin and eosin for 15 min and 1 min respectively and dehydrated in xylene as previously indicated^65^. For immunofluorescence staining, cross sections of muscle tissues (TA, EDL, SOL) were fixed in 4% PFA for 15 mins and quenched with 100 mM glycine for 10 min. Fixed tissue sections were then incubated with Blocking Buffer (5 % goat serum, 2 % bovine serum albumin, 0.1 % Triton X-100, and 0.1 % sodium azide, 1X PBS) for 2 hr. Tissue sections were incubated with primary antibodies diluted in blocking buffer overnight at 4°C and incubated with secondary antibodies and DAPI for 1 hr at room temperature. For Oil Red O (ORO) staining, muscle tissues were fixed in 4 % PFA for 30 min and stained using Oil red O working solutions for 60 min. After washing with running water for 2 min, it was counterstained with hemoxylin, and imaged. All images are representative results of at least four biological replicates. For Succinate dehydrogenase (SDH) staining, muscle tissues were stained with SDH solution (10ml 0.2 M phosphate buffer, 270 mg sodium succinate, 10 mg NBT) for 10-15 min. The slides were rinsed with deionized H_2_O and mount the coverslips with mounting medium.

### Treadmill test and grip strength measurement

Treadmill exercise testing was performed as previously described^66^. WT, *Prmt5^MKO^* and *Prmt5/Pnpla2^MKO^* mice (2 months old), were placed on a treadmill (Eco3/6; Columbus Instruments, Columbus, OH) for 10 min for 3 consecutive days at constant 10 m/minute speed for acclimation. On the testing day, mouse ran on the treadmill at 10 m/minute for 5 min and the speed was then increased by 2 m/minute every 2 min until mice were exhausted. Exercise capacity, including running distance, running time and maximum speed was measured. For grip strength, WT, *Prmt5^MKO^* and *Prmt5/Pnpla2^MKO^* mice (2 months old) were tested to measure using a DFE II series Digital Force Gauge (Ametek DFE II 2-LBF 10-N) with an attached metal grid. The mouse was allowed to grasp the metal grid an, and the mouse tail was gently pulled along the axis of the grid. The peak tension at the time of release was recorded. The grip strength was measured three times and the average strength was normalized to body weight (N/g)

### Co-immunoprecipitation, protein extraction and western blot analysis

For Co-IP, C2C12 cells were transfected with pcDNA-GFP-PRMT5 and pcDNA3.1-2×FLAG-SREBP1a, and cells were harvested after 24 hr. 500 μg protein lysate was precleared with protein A/G agarose bead at 4 °C for 2 hr, and 1 ug of primary anti-FLAG, or anti-GFP was added into the protein lysate, and rotate for 4 hours, followed by addition of protein A/G agarose bead and rotate for overnight.

Total protein was extracted from homogenized muscle tissue using RIPA buffer containing 25 mM Tris-HCl (pH 8.0), 150 mM NaCl, 1 mM EDTA, 0.5 % NP-40, and 0.1 % SDS) supplemented with Proteinase inhibitor and phenylmethylsulphonyl fluoride (PMSF). The concentration of supernatant proteins was quantified using Peirce BCA Protein Assay Reagent (Pierce Biotechnology). Proteins were separated by electrophoresis, transferred to polyvinylidene fluoride (PVDF) membrane, blocked with 5 % fat-free milk for 1 hr at room temperature and incubated with primary antibodies overnight at 4 °C. Antibodies used for western blot analysis were listed in Supplementary Table 2. Immunodetection was detected using enhanced chemiluminescence western blotting substrate (Santa Cruz Biotechnology) on a FluorChem R system (Proteinsimple).

### Chromatin immunoprecipitation

C2C12 cell-derived myotubes and muscle tissue from both WT and *Prmt5^MKO^* mice were crosslinked with 1 % formaldehyde for 10 min at room temperature and quenched by the addition of 125 mM glycine for 5 min at room temperature. After the samples were washed cold PBS, they were incubated with ChIP cell lysis buffer (20 mM Tris pH 8.0, 0.1%, 85 mM KCl, 0.5% NP40) supplemented with protease inhibitor. After centrifugation, the nuclei were resuspended in nuclei lysis buffer (50 mM Tris pH 8.0, 10 mM EDTA, 1% SDS) and sonicated. The supernatant was used for immunoprecipitation with the indicated antibodies (PRMT5, H4R3Me2s). The immunoprecipitants were eluted and reverse crosslinked overnight at 65 °C. Phenol-Chloroform method was used to purify DNA fragments and qRT-PCR was performed as indicated. A list of primers used is provided in Supplementary Table 1. Results were presented as mean ± s.d. from three independent experiments.

### Assessment of muscle contractile function

Contractile properties of the slow-twitch soleus and the fast-twitch EDL muscles were measured by using an *in vitro* muscle test system (1200A Intact Muscle Test System, Aurora Scientific). Briefly, the hindlimb was excised under isoflurane anesthesia, and placed in a bicarbonate-buffered solution (137 mM NaCl, 5 mM KCl, 1mM MgSO_4_, 1 mM NaH_2_PO_4_, 24 mM NaHCO_3_, and 2 mM CaCl_2_) equilibrated with 95% O_2_-5% CO_2_. The assessment of contractile function of the EDL muscle was performed first followed by the Soleus muscle. After surgical isolation, braided silk suture thread (4-0, Fine Science Tools) was tied around each end of the muscle tendons. The muscles were then transferred to a tissue bath apparatus containing a bicarbonate-buffered solution at room temperature continuously bubbled with carbogen (95% O_2_-5%CO_2_) and mounted between two platinum electrodes (1200A Intact Muscle Test System; Aurora Scientific). After the optimal muscle length was determined, the temperature of the glass bath was increased to 32°C, and the muscles were thermally equilibrated for 10 min. The force-frequency relationship was then generated by selected frequencies between 1-300 Hz for the EDL muscle, and 1 −200 Hz for Soleus muscle. After completion of the assessment, the muscle was removed from the organ bath, trimmed of connective tissue, blotted dry, and weighed. Muscle cross-sectional area (CSA) was determined by dividing the wet muscle mass by the product of Lo and muscle-specific density (1.056 g/cm^3^). Specific force (N/cm^2^) was calculated by dividing the muscle force (N) by the CSA (cm^2^)

### Whole body composition analysis

The Echo-MRI 130 analyzer (EchoMRI LLC, Houston, TX, USA) was used to measure body composition of live mice. After the calibrating the instrument with corn oil, the mouse was gently placed into a cylindrical holder. The holder was inserted into the EcoMRI^™^ system to measure lean mass, fat mass, and water weight.

### Total RNA extraction and qRT-PCR

Total RNA was extracted from cells and tissues using TRIzol reagent (Thermo Fisher Scientific) according to the manufacturer’s instruction. 2 μg of total RNA were reversed transcribed with random primers, M-MLV reverse transcriptase and DTT. Real-time qPCR was carried out in a Roche Light cycler 480 PCR system with SYBR green master mix and gene-specific primers, listed in Supplementary Table 1. Relative changes in gene expression were analyzed using the 2^-ΔΔCT^ method and normalized to β-actin.

### Seahorse OCR measurement

Primary myoblasts cells (1×10^5^ cells) isolated from WT and *Prmt5^MKO^* mice were seeded in Matrigel coated XF24 microplates (SeaHorse, bioscience). After differentiation for 3 days, myotubes were washed three times with XF medium (supplemented with SeaHorse XF RPMI medium, 5 mM glucose, 2 mM pyruvate, 1 mM glutamine, pH 7.4), and pre-incubated in XF medium for 1 hour at 37 °C in a non-CO_2_ incubator. Oligomycin (3 μM), FCCP (3 μM), Rotenone (1.5 μM), Antimycin A (1.5 μM) were preloaded into cartridges and injected sequentially into XF wells to monitor oxygen consumption rate, OCR (pmol/min). All mitochondria respiration rates were calculated by the SeaHorse Wave software and normalized to the cellular protein contents.

### Statistical Analysis

All data are presented as mean ± standard error of the mean (S.E.M). All quantitative analyses were conducted with Student’s *t*-test and a two-tail distribution calculated using the GraphPad Prism. Comparisons with *p* values < 0.05 were considered statistically significant.

## Data availability

The data that support the findings of this study are available from the corresponding author on request.

## Acknowledgements

This work was supported by grants from the US National Institutes of Health (R01AR071649) to S.K. We thank Dr. Changdeng Hu (Purdue University, USA) for kindly providing the *Prmt5^flox/flox^* mice, and Dr. Bruno T. Roseguini for technical assistance for contractile muscle force measurement. We also thank Jun Wu, and Mary Larimore for mouse colony maintenance and technical support, and members of the Kuang Laboratory for critical comments.

## Author Contributions

K.H.K., and S.K. conceived the project, designed the experiments, and prepared the manuscript. K.H.K., Z.J., M.S., J.C., J.Q., S.O., X.C., S.S., and F.Y. performed experiments and analyzed the data. C.H., B.T.R., and A.N.I. provided reagents.

